# Computational Analysis of the Metal Selectivity of Matrix Metalloproteinase 8

**DOI:** 10.1101/2020.06.30.165720

**Authors:** Zheng Long

## Abstract

Matrix metalloproteinase (MMP) is a class of metalloenzyme that cleaves peptide bonds in extracellular matrices. Their functions are important in both health and disease of animals. Here using quantum mechanics simulations of the MMP8 protein, the coordination chemistry of different metal cofactors is examined. Comparisons found that Jhan-Teller effects in Cu(II) destabilize the wild-type MMP8 but a histidine to glutamine mutation at residue number 197 can potentially allow the MMP8 protein to utilize Cu(II) in reactions. Simulations also demonstrates the requirement of a conformational change in the ligand before enzymatic cleavage. The insights provided in here will assist future protein engineering efforts utilizing the MMP8 protein.

## Introduction

Matrix metalloproteinase (MMP) is a class of proteins whose native functions involves the processing of extracellular matrix and cytokines.^1-2^ As a result of their involvement in cytokine cleavage, they are also part of the signal transduction pathway involving immune cells. MMPs have been used in anti-cancer clinical trials^3^, however, their functions and involvements in cancer has not been fully understood. While still believed to be viable targets in cancer therapies, targeting of individual MMPs is believed to be important for the success of the approach ^4^. The understanding of MMP protein metal selectivity can help to elucidate the behaviors of these proteins in healthy tissues as well as disease environments. The aim of this work is to use computational methods to provide insights into the metal selectivity of the MMPs to assist in future efforts to engineer MMPs that may serve therapeutic functions.

All MMPs contain three important domains, the pro-domain, catalytic domain and hemopexin like domain. The catalytic domain of MMPs are sufficient for their enzymatic cleavage^1^. It is known that Zn(II) bound to MMPs catalytic domain can be substituted by Cu(II), Co(II), Mg(II), and Mn(II) with various levels of activities that are concentration dependent. In addition, metal depletion is a reversible process where replenishment of the metals revives protein activities ^5^. These results suggest that the MMP catalytic chemistry is not unique to Zn(II) ions. However, different metal cofactors demonstrate different levels of activities and the previously published results are tabulated in Table 1 this data shows significant bias in MMPs against metal cofactors. In experimental data, Mg(II) has slightly lower activities compared to Zn(II), in contrast, Cu(II) demonstrates almost no activity at the concentration of 10 mM. However, 10 mM is still higher than physiologically relevant metal concentration in the body ^6-7^, as a result, it is possible that at even lower concentrations, more differences can be observed in MMPs.

**Table 1.**
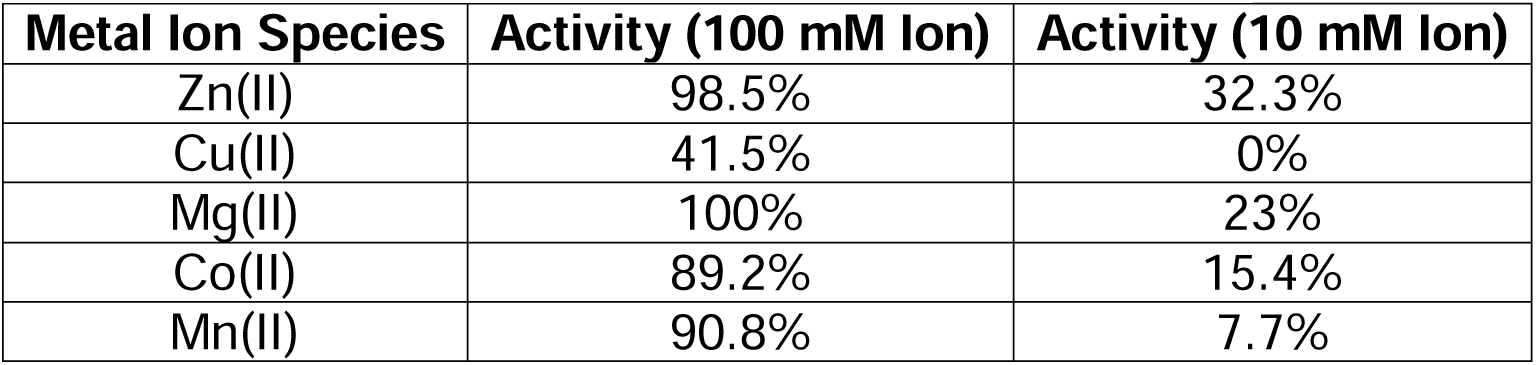
results for metal activities against collagen ^5^.

| **Metal Ion Species** | **Activity (100 mM Ion)** | **Activity (10 mM Ion)** |
| --- | --- | --- |
| Zn(II) | 98.5% | 32.3% |
| Cu(II) | 41.5% | 0% |
| Mg(II) | 100% | 23% |
| Co(II) | 89.2% | 15.4% |
| Mn(II) | 90.8% | 7.7% |

In MMPs, selectivity of divalent metals is not evident from the coordination site structure, for similar coordination chemistry can be found in proteins harboring different divalent metal ions ^8-9^. Experimental data suggests that there is physiological relevance of MMP proteins using Cu(II) as a cofactor, this is despite Cu(II) MMP seem to demonstrate no collagenase activities under normal conditions. In fact, it was shown that MMPs can process new ligands in the presence of Cu(II) ^10-12^, this suggest that changing metal cofactors in MMPs alters ligand selectivity. Interestingly, there exist a variant metalloproteinase favoring Cu(II) instead of Zn(II), which can serve as a comparison in the simulations with MMPs.

The discovery and characterization of a new family of metalloproteinase in the green algea *Volvox carteri* named the VMP3 reveal a similar metal coordination motif of QEXXHXXGXXH instead of the HEXXHXXGXXH in the MMPs, and the discovery that this motif favors the utilization of copper in catalysis ^13^ suggests that a similar mutation in the MMPs can alter Zn(II) and Cu(II) selectivity in the MMP catalytic site metal binding pocket. Comparing the coordination chemistry of the two proteins might reveal the mechanism behind metal selectivity of MMPs and VMP3. However, there are currently no structures available for VMP3, thus a homology model was generated for the apo VMP3 using IntFold ^14^. The structure of the predicted VMP3 protein is shown in Figure 1B. The QEXXHXXGXXH motif is shown arranged spatially similar to that of MMP8 (Figure 1A). Compared to VMP proteins, MMP proteins has been studied extensively, and many structures exist for MMPs.

**Figure 1.**
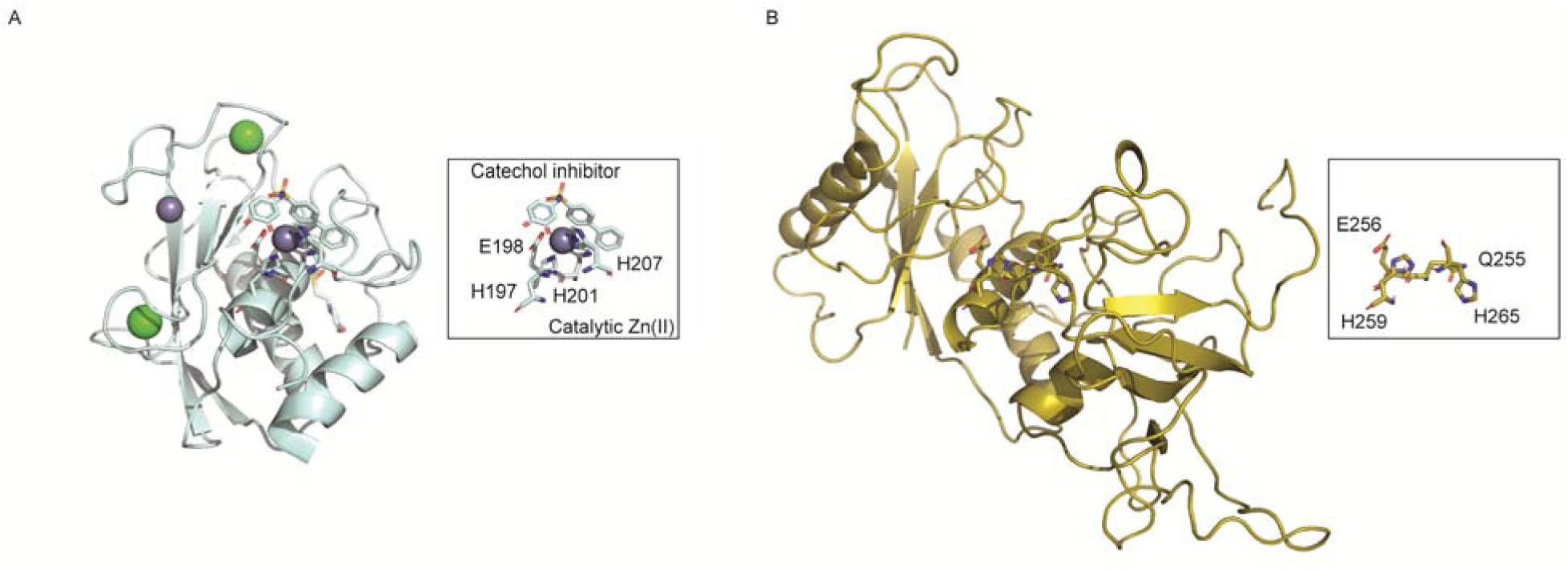
(A) Inhibited form of the MMP8 catalytic domain crystal structure (5h8x) ^19^, the rectangular box highlights the catalytic site in MMP8 with the Zn(II) ion. (B) Homology modeled apo VMP3 structure predicted using IntFold ^14^. The rectangular box highlights the catalytic site in VMP3 without the Zn(II) ion.

In humans, there are at least 10 MMP proteins: MMP1, MMP2, MMP3, MMP7, MMP8, MMP9, MMP10, MMP11, MMP20, and MMP26. MMP8 or neutrophil metalloproteinase is expressed on neutrophils to remodel the extracellular matrix allowing immune cell infiltrate tissues. It is believed to possess some anti-cancer effects by allowing immune cell penetration into the cancer microenvironments. MMP8 itself is involved in tumor survival and mobility, its expression is associated with both pro and anti-tumor effects in different types of tumor environments ^15^. Understandings of the functional metal coordination of the MMP catalytic sites can help to elucidate the effects of divalent metals as well as mutations over the ligand selectivity of the MMP proteins in different disease environments. Because of the importance of MMP8 in disease, it is used as a representative MMP protein for simulations. In addition, due to the disease relevance of Cu(II) in cancer where it is found that the concentration of Cu(II) is elevated in tumor tissues ^7, 16-17^, focus has also been given to the effects of Cu(II) cofactor on the structures of the MMP8 protein.

The MMP8 protein contains two Zn(II) binding sites. Site I has four residues of H147, D149, H162, and H175 forming a tetrahedral coordination site with the zinc ion. Site II has three residues H197, H201, and H207 forming a zinc binding site with the motif HEXXHXXGXXH. Site II is the catalytic site in MMP8 where peptide bond cleavage takes place. The motif in site II is not just found in other MMPs but also common to a family of enzymes named metzinsins ^18^. Because of the conserved catalytic site structures across different MMPs, the chemistry of MMPs can be readily modeled base on the coordination sites around the metal center, and the results can be readily applied to other proteins within the class.

Close examination of the crystal structure of inhibitor bound MMP8 reveals that the zinc ion in the catalytic site is coordinated by three histidine residues and one catechol inhibitor N-(3,4-dihydroxyphenyl)-4-diphenylsulfonamide ^19^. This crystal structure of MMP8 is shown in Figure 1A. In the coordination center highlighted in the boxed area, Zn(II) is stabilized by the addition of coordinating inhibitors. In the absence of the inhibitors or ligands, the coordination site is believed to be satisfied by a water molecule which can be polarized by the adjacent glutamate residue next to one of the coordinating histidine residues. This has been demonstrated in the inhibitor-free MMP-12 structure ^20^ and in MMP-3 ^21-23^ previously. These knowledge forms the basis of the modeling performed in this work on MMP8, because the structures of the catalytic sites of all MMPs are conserved, metal coordination chemistry is believed to be conserved for all MMPs.

MMP ligand interactions has been studied using x-ray crystallography and nuclear magnetic resonance spectroscopy ^24-25^. Using these structural information, the direction of the ligand-MMP interactions can be identified, also previous simulations has suggested that the ligand coordination likely occurs in a bi-dentate fashion.^21^ These prior knowledge enables the correct modeling of the ligand structure at the MMP8 catalytic site.

There is no crystal structure for copper bound MMP proteins but similar copper proteins exist. Plastocyanins contain a type I copper center ^26^ which is formed by two histidine residues, one cysteine, and one methionine into a distorted tetrahedral coordination center around the Cu(II) ion. The three histidine coordination for Cu(II) typically has pyramidal coordination involving a second Cu(II) nearby bonding to each other through the tips (type III copper center)^26^. For proteins like plastocyanin, metal requirements are also non-exclusive ^27^. In azurins for example, Zn(II) occupied active site instead of the native Cu(II) remains functional, however, a geometry distortion was observed in the Zn(II) occupied protein ^28^. These results suggest that in proteins utilizing divalent metals as cofactors, the metal species are often times non-exclusive but the presence of non-native metal ions is still disruptive to the coordination geometry.

Using computational simulations, it is possible to understand the behaviors of MMPs and make predictions of effects due to changes in the coordination centers. These approaches have been instrumental for studies of metal coordination chemistry within proteins. Metal coordination chemistry requires the use of quantum mechanics (QM) simulations, for the traditional force fields have not been able to completely address the chemistry around these metal co-factors.^29^ Previous study has identified the presence/absence of metal binding selectivity in various type of divalent metal metalloproteins using molecular dynamics simulations, this approach, however, does not provide enough accuracy in comparisons of structural differences induced by different divalent metal cofactors ^30^. Density functional theory (DFT) has been applied to the studies of protein, which provides sufficient computational efficiency as well as accuracy. But compared to molecular dynamics simulations, only part of the protein can be simulated efficiently and thus this general approach is applied here to only simulate around the first coordination shell around the metal coordination center.^31^ The QM approach undertaken here is beneficial for the close examination of the local coordination chemistry, however, it largely misses the other protein-protein interactions important for ligand selectivity as well as complex stability, thus the scope of this work is limited to the events around the coordination center, and a more sophisticated approach using the QM/MM methods has to be utilized to understand the behavior of the MMPs systematically ^32^.

## Methods

Catalytic site modeling is performed in PyMol. Computations were performed with a Linux system equipped with a NVidia Tesla K10 GPU. DFT Geometry optimization were performed with BLYP3/LANL2DZ level of theory as done previously ^21^ in Gaussian 09 (www.gaussian.com). Energy calculations were performed with B3PW91/6-311G++3df,3dp level of theory. Canonical molecular orbitals calculated with the same levels of theory were visualized in GaussView 5.0 (www.gaussian.com) using B3PW91/6-311G++3df,3dp methods.

Structural modeling was carried out starting from the Zn(II) catalytic site for MMP8 in crystal structure ^19^ and the position of the water molecule was modeled in before geometry optimization based on the previous structures and structural models ^20, 25^. At the edge of the computational boundary, the Cα atoms of the amino acid residues are replaced by methyl groups, and the carbon atom is frozen in calculation to prevent forbidden backbone angles without the contextual restraints. This approach is chosen based on the observation that there are no separate conformations for apo and ligand bound/inhibited form of the MMP protein ^19, 25^. Ligand modeling was based on the crystal structure of the collagen III derived peptides ^33^ which contain known cleavage sites for MMP8 and structural models of MMP collagen interactions ^25^.

## Results and Discussion

Because of the differential activities of different metal ions, computational methods were used to elucidate the molecular mechanism of the metal biases. First, coordination geometries of different metals in the catalytic site was examined after performing a series of geometry optimizations starting from the protein crystal structure containing the Zn(II) cofactor (Figure2). Two major types of coordination geometries can be identified in the simulations for the MMP8 catalytic site in the absence of E198 or ligands. Co(II), Cu(II) and Ni(II) coordination geometries are similar to square planner while Ca(II), Zn(II), Mg(II) and Mo(II) geometries are similar to tetrahedral. Mn(II) coordination geometry, however, resides between the two cases.

In the calculations, the catalytic site was simplified such that calculations are efficient and cluster structure is representative of the coordination chemistry around the metal center. In the modeling, the alpha carbons were converted to methyl groups to prevent unrealistic destabilization at the simulation boundaries. In addition, restraints were added to the system to prevent the residues from deviating from their natural orientations. The observation that different metal ions all have distinctive geometries in the simulations is expected and suggests ligand selectivity is dictated to some levels by the coordination chemistry of the metals and such differences might be correlated with the functional differences demonstrated by these metals. Geometric differences around the coordination sites are observed while the C_α_atoms of the amino acid residues are fixed, indicating that these rearrangements can be accommodated by local side-chain conformational rearrangements and thus does not necessarily require backbone conformational changes. These fixed atoms are also important for they can prevent unrealistic geometries which may require sterically forbidden backbone conformations, and it is especially important for subsequent calculations where additional residues are included which further complicates the energy landscape.

In the ligand-free protein, the effects of the H197Q mutation is examined to determine if the mutation is structurally viable. Simulations show that this mutation can fit within the catalytic site of MMP8 without requiring protein backbone changes (Figure 3). Simulations of the H197Q mutants also show that coordination geometries around Cu(II) and Zn(II) are changed due to changed coordinating chemical groups. In Figure 3 it can be seen that, compare to the wild-type protein (Figure 2 A-D), H197Q mutant has distortion of the coordination geometry of the Zn(II) or Cu(II) cofactors mostly due to the way Q197 approaches the metal center. These differences are consistent with observations when ligand is included in the simulations.

**Figure 2.**
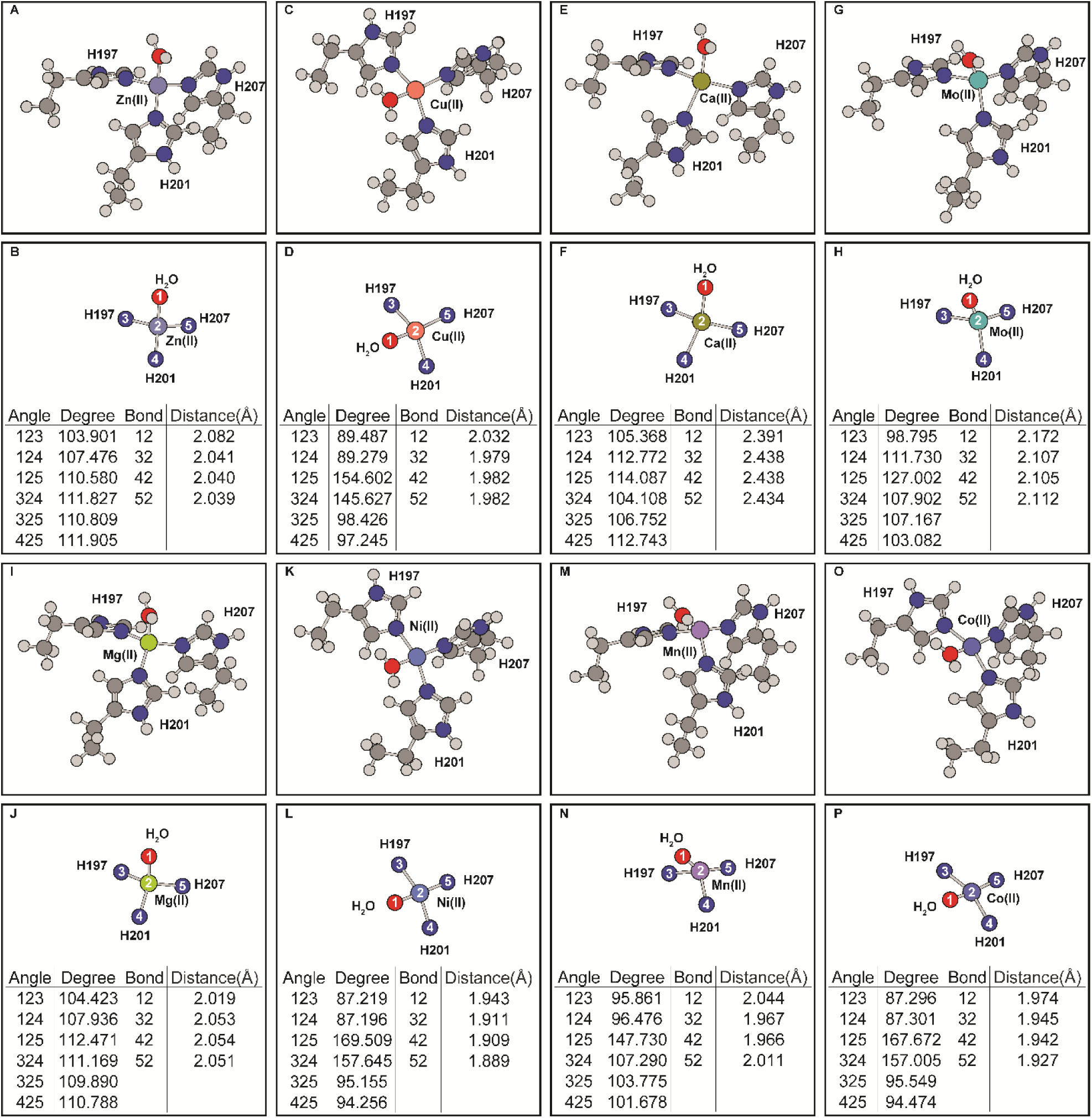
Effects of divalent metal occupancy on the MMP8 catalytic center without E198 or ligand. (A-B) Zn(II). (C-D) Cu(II). (E-F) Ca(II). (G-H) Mo(II). (I-J) Mg(II). (K-L) Ni(II). (M-N) Mn(II). (O-P) Co(II). (A), (C), (E), (G), (I), (K), (M), and (O) are the full coordination shell. (B), (D), (F), (H), (J), (L), (N) and (P) contain simplified depictions showing only atoms directly coordinated with the metals, as well as tables for the angles and distances of the coordination center.

**Figure 3.**
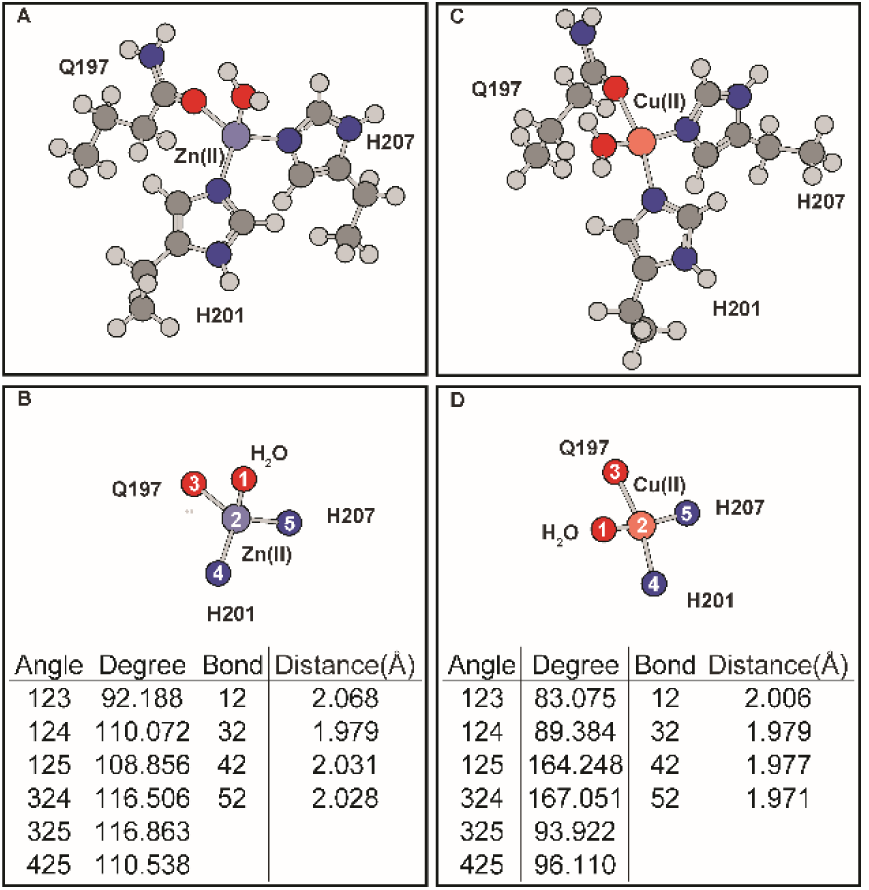
Effects of H197Q mutation on the coordination geometry of the MMP8 catalytic site without E198 or ligand. (A-B) Zn(II). (C-D) Cu(II). (A) and (C) are the full coordination shell. (B) and (D) contain simplified depictions showing only atoms directly coordinated with the metals, as well as tables for the angles and distances of the coordination center.

Inclusion of E198 in the calculations distorts both the wild-type and mutant coordination geometries (Figure 4). In wild-type protein, E198 side-chain draws the water molecule closer to itself, thus changes the coordination angles (Figure 4 AB and CD). Similar effects are seen in H197Q mutant with Cu(II) center (Figure 4 GH). The distortions caused by E198 reduces the coordination geometries differences between wild-type or H197Q mutant catalytic site. However, after including E198 in simulations, the coordination geometries of Zn(II) occupied MMP8 H197Q catalytic site produces an additional hydrogen bond formed between E198 carbonyl oxygen atom and Q197 side-chain ε amino group (Figure 4E and 4F). This hydrogen bond changes the relative positions of residue H207 to residue 197. In contrast, hydrogen bonding interactions are absent when Cu(II) is present in the mutant catalytic site (Figure 4G and 4H). Such inter-residue hydrogen bonding interactions are also absent in wild-type MMP8 catalytic site with Cu(II) or Zn(II) occupancy.

**Figure 4.**
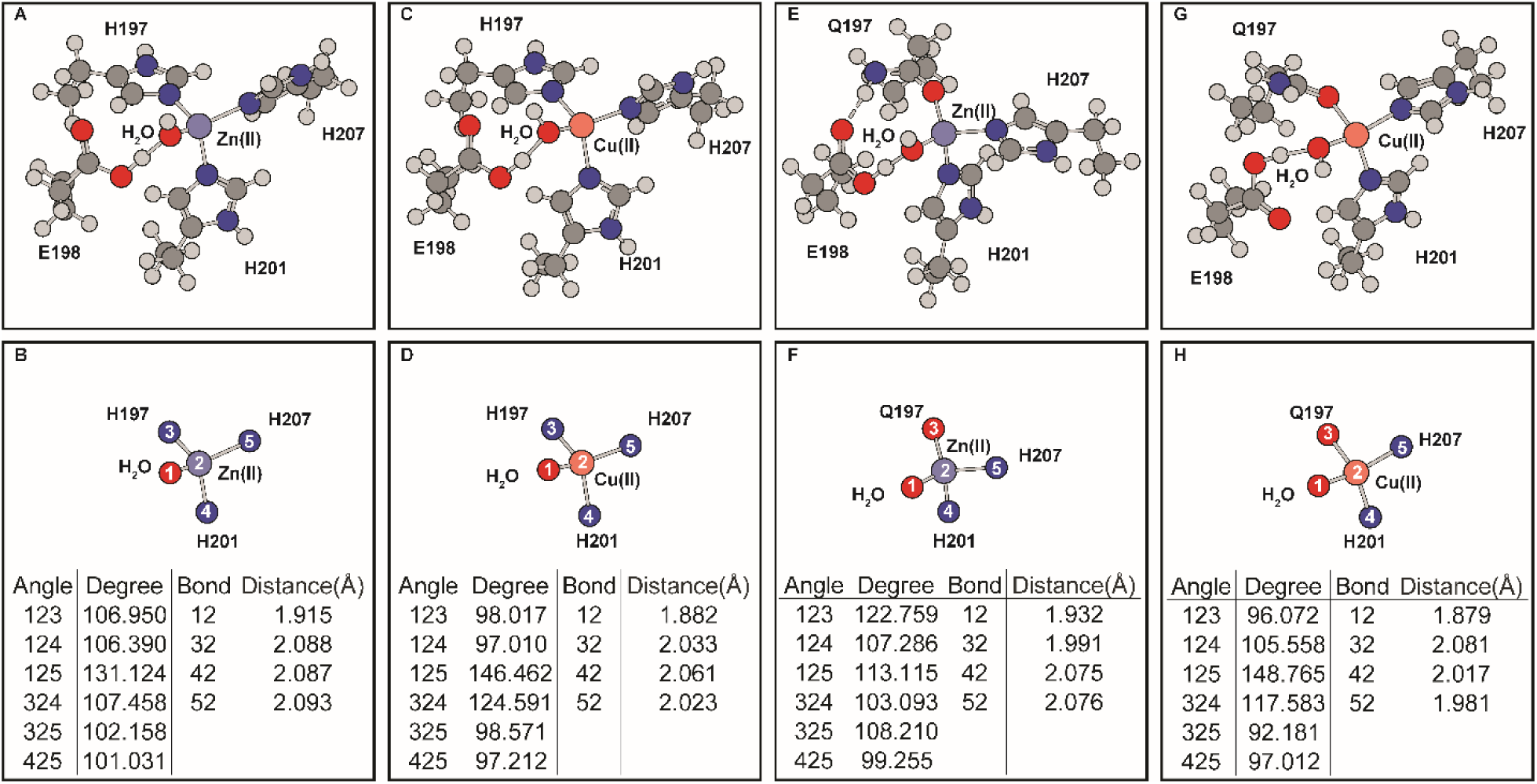
Effects of E198 on the MMP8 catalytic center without ligand. (A-B) Zn(II) occupied wild-type MMP8. (C-D) Cu(II) occupied wild-type MMP8. (E-F) Zn(II) occupied MMP8 H197Q mutant. (G-H) Cu(II) occupied MMP8 H197Q mutant. (A), (C), (E), and (G) are the full coordination shell. (B), (D), (F), and (H) contain simplified depictions showing only atoms directly coordinated with the metals, as well as tables for the angles and distances of the coordination center.

In addition to the inclusion of E198, ligand binding alters the coordination geometry further (Figure 5A-D). The coordination of the ligand in the native Zn(II) occupied MMP8 catalytic site forms a distorted octahedral geometry which involves bidentate coordination from two adjacent ligand carbonyl groups. From the two types of coordination geometries observed in the ligand-free simulations (Figure 2), additional metal ions Mg(II) and Co(II) (Figure 5E-H) were selected as comparisons to Zn(II) and Cu(II) (Figure 5A-D). Results show that similar to the ligand-free simulations, the coordination geometry of Mg(II)-MMP-ligand complex is similar to that of the Zn(II)-MMP8-ligand geometry while Co(II)-MMP8-ligand geometry resembling that of the Cu(II)-MMP8-ligand geometry. From the geometry of the Cu(II)-MMP8-ligand structure, the coordination of the ligand is somewhat incompatible with the coordination site, because the i-1 ligand does not seem to be able to get close to the metal. In the Cu(II) occupied wild-type MMP8 catalytic site, the coordination bond between Cu(II) and carbonyl oxygen atom of the ligand at i position extends to 3.35 Å which is significantly larger than the regular coordination bonds in the range of 2.00 −2.50 Å distance range measured in structures containing other metal ions.

**Figure 5.**
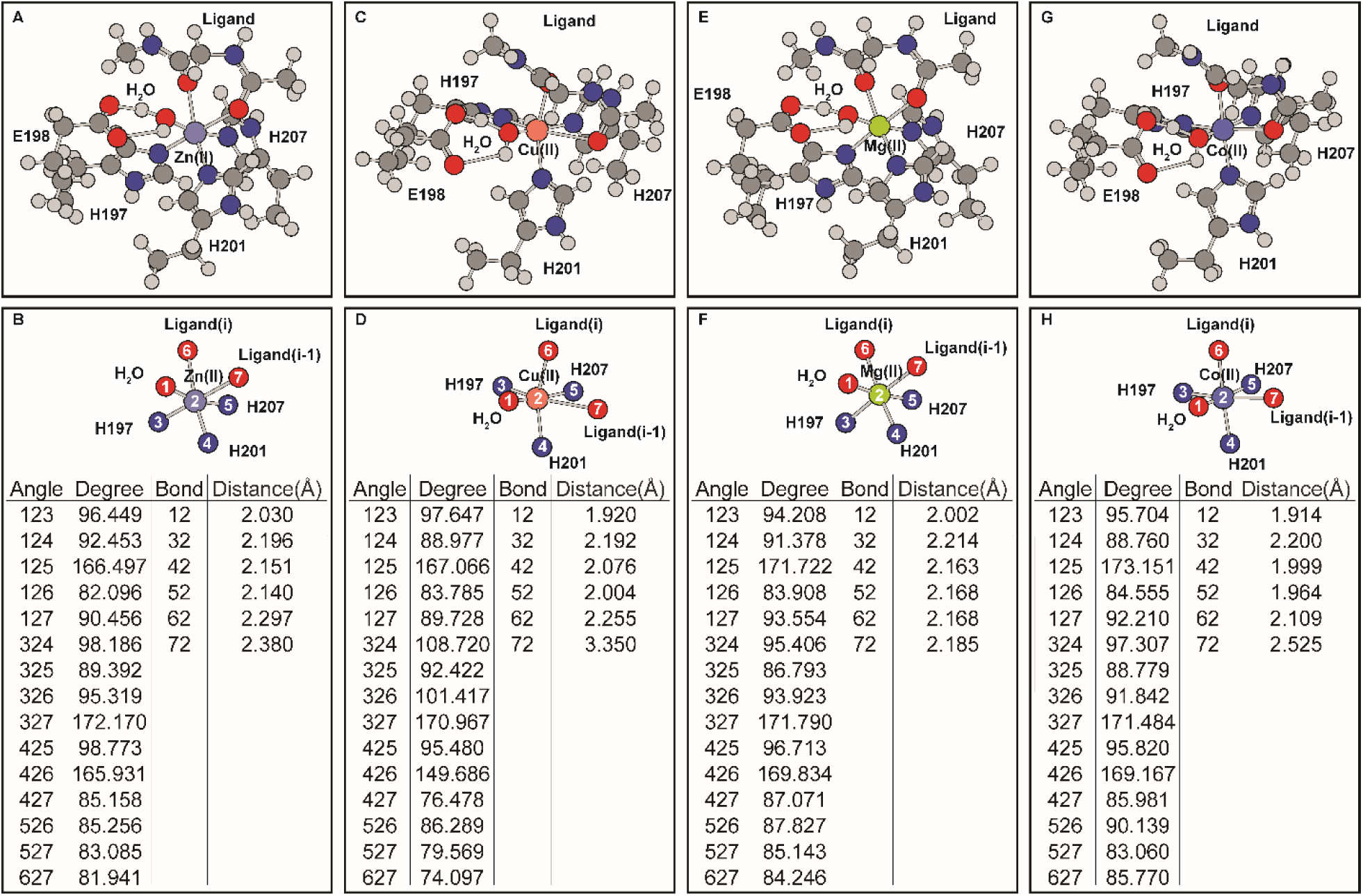
Effects of divalent metal occupency on the ligand bound MMP8 catalytic center. (A-B) Zn(II). (C-D) Cu(II). (E-F) Mg(II). (G-H) Co(II). (A), (C), (E), and (G) are the full coordination shell. (B), (D), (F), and (H) contain simplified depictions showing only atoms directly coordinated with the metals, as well as tables for the angles and distances of the coordination center.

The differences in coordination geometry due to histidine to glutamine mutation is expected because the two amino acids contain sidechains that are very different in coordination properties. Strong Jahn-Teller effect, indicated by the distortion of Cu(II) – ligand_i_ bond and the elongation of the Cu(II) – ligand_i-1_ bond (Figure 5C-D) is observed in Cu(II) occupied wild-type MMP8 active site. Similar Jahn-Teller effect is observed in Co(II) occupied wild-type MMP catalytic site although to a lesser extent. Jahn-Teller effect in Cu(II) occupied H197Q MMP8 mutant (Figure 6C-D) on the other hand is reflected in the elongation of the Cu(II) – ligand_i_ and Cu(II) – H201 bonds. Distortion is also observed in Zn(II) occupied H197Q mutant in the presence of the ligand, this is due to the formation of hydrogen bond between the side-chain NH_2_ group with the ligand carbonyl oxygen at position i. These distortions constitute the metal selectivity of the MMP protein catalytic site by changing the ligand-MMP complex stabilities.

**Figure 6.**
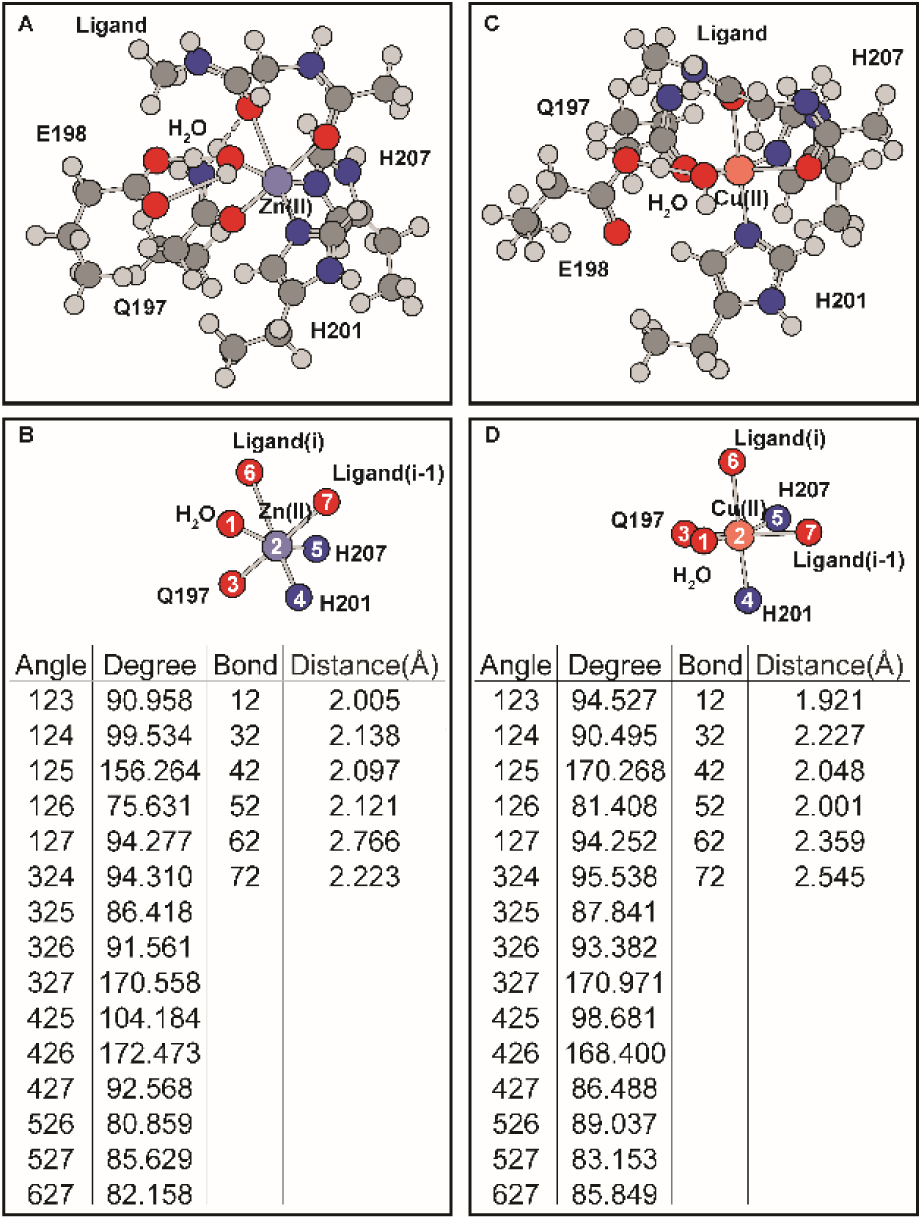
Effects of H197Q mutations on the ligand bound MMP8 catalytic center. (A-B) Zn(II). (C-D) Cu(II). (A) and (C) are the full coordination shell. (B) and (D) contain simplified depictions showing only atoms directly coordinated with the metals, as well as tables for the angles and distances of the coordination center.

The consequences of the distortions in the Cu(II) occupied protein is significant. Compared to the H197Q mutant, wild-type catalytic site contains ligand coordination bond that stretches to 3.35 Å. This prevents the orbital hybridization between the orbitals of the Cu(II) ion with the orbitals of the carbonyl oxygen atom. This is demonstrated in Figure S1 where the molecular orbitals containing all the bonding orbitals between the Cu(II) and ligand atoms are shown, Cu(II) in wild-type MMP8 catalytic site has no bonding electron density between Cu(II) and the carbonyl oxygen atom at ligand position i-1. As a result, this complex is less stable. Thus, it can be concluded here that the Jahn-Teller distortions in Cu(II) bound MMP8 destabilizes the coordination center. The effect of the hydrogen bonding in Zn(II) occupied H197Q mutant also elongates the metal – ligand bond to 2.77 Å which still allows orbital hybridization revealed by similar molecular orbital analysis, although the bonding molecular orbitals reduces from twelve in wild-type MMP8 to seven in the H197Q mutant. As a result, conclusion cannot be drawn about the effect of the mutation on the stability of the Zn(II) bound MMP8 H197Q.

Close examination of the distances between the water oxygen atom and the ligand carbonyl carbon atom at ligand position i where cleavage takes place, shows that this distance is around 2.8-2.9 Å in all of the calculated structures (Table S1). Mutations and metal induced changes does not change this distance significantly. As a result, no implications can be made to the reaction rate based on our simulations. Furthermore, due to the lack of information about the mechanisms of the induced transition state, i.e. whether it is due to dynamic changes in the MMP protein or due to changes in ligand orientation, a confident prediction of the transition state structure was not sought here using simulations.

Compared to the functional data shown in Table 1 which shows that the activities of Mg(II) is slightly lowered than Zn(II) at 10 mM concentration, our simulation shows Mg(II) demonstrates very similar coordination geometry in MMP8. In contrast, Cu(II) has no activities against collagen at 10 mM concentration. Our simulation shows that Cu(II) destabilizes the ligand-metal-MMP8 complex. In experiments, Co(II) has higher activities than Cu(II) against collagen but much lower activities than Mg(II) or Zn(II), this is compared to our simulations where the Co(II) bound MMP8 experiences less distortion effect compared to Cu(II) bound protein.

Active site mutants of the MMP protein has been made experimentally. The mutation in the active site motif from HEXXHXXGXXH to HQXXHXXGXXH has been studied. This is a loss-of-function mutation that diminishes the catalytic activities of the MMP proteins, and the structure of the mutant is comparable to that of the wild-type protein.^34^ In contrast experimental results for QEXXHXXGXXH mutations does not exist, so functional comparisons can only be made with the VMP3 protein which harbors a naturally functional QEXXHXXGXXH motif favoring Cu(II). The observation that in the H197Q mutant, which is analogues to VMP3, Cu(II) induces a more stable coordination structure than in wild-type MMP8 by allowing an additional ligand coordination bond suggest that the H197Q might demonstrate similar metal selectivity to VMP3 where Cu(II) will have much better utilization.

The computations also suggest that favorable ligand torsion angles for metal coordination deviates from that measured in collagen crystal structures. These torsion angle changes are measured and shown in Table S2 showing comparisons of the angles found in geometry optimized structures to the starting structure of collagen III. Previous studies hypothesize that partial refolding of the collagen might occur leading to collagen cleavage ^24^, this matches previous results showing that the torsion angles of the bound ligand changes significantly. Ligand binding to the active site metal can potentially induce structural changes in MMPs as well. However, based on the simulations, large domain conformational changes might not be required, for side-chain rearrangements can be sufficient to accommodate these local changes.

Evidence suggest Cu(II) does lead to additional substrate proteolysis in MMPs compared to Zn(II) ^11^ these effects are reflected in the selectivity of torsion angles for different ligands, limited modeling capabilities of the QM simulations likely means that simulating larger segments of the MMP proteins complexed with a more contextual ligand will remain difficult. This can potentially be overcome using novel potential energy functions in MM simulations ^35^.

**Figure S1.**
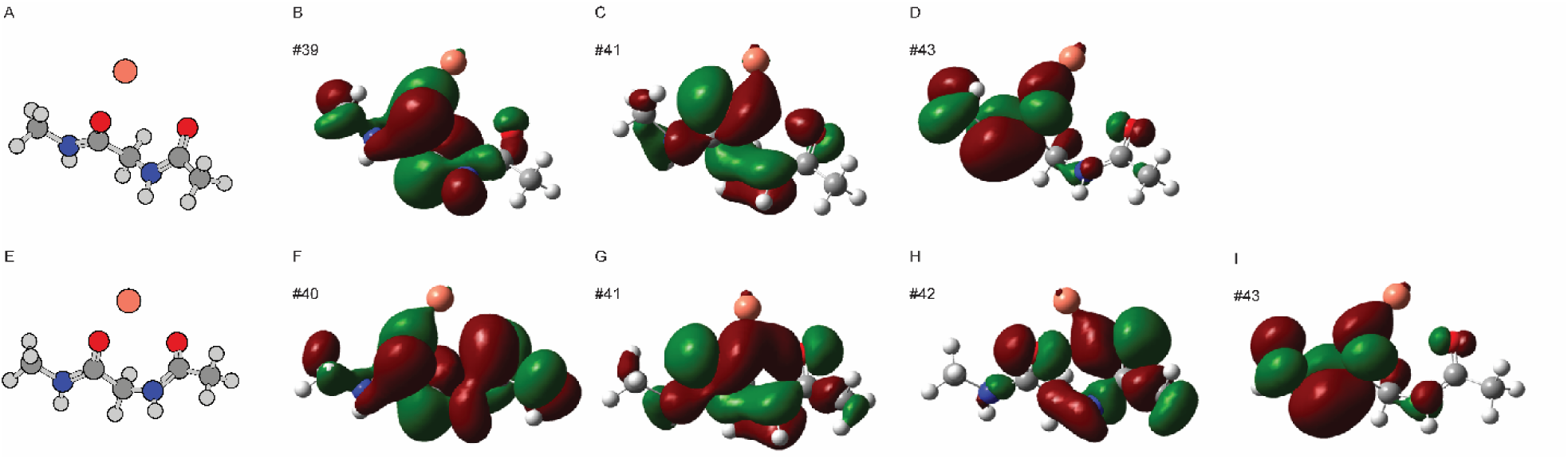
Cu(II) –ligand orbital mixing the bonding orbitals formed between the ligand and Cu(II) ion are shown. (A) is a molecular representation of the Cu(II)-ligand complex in wild-type MMP8 catalytic site. (B)-(D) are three bonding molecular orbitals between Cu(II) and ligand wild-type MMP8 catalytic site. (E) is a molecular representation of the Cu(II)-ligand complex in H197Q mutant catalytic site. (B)-(D) are four bonding molecular orbitals between Cu(II) and ligand in H197Q mutant catalytic site.

**Table S1.**
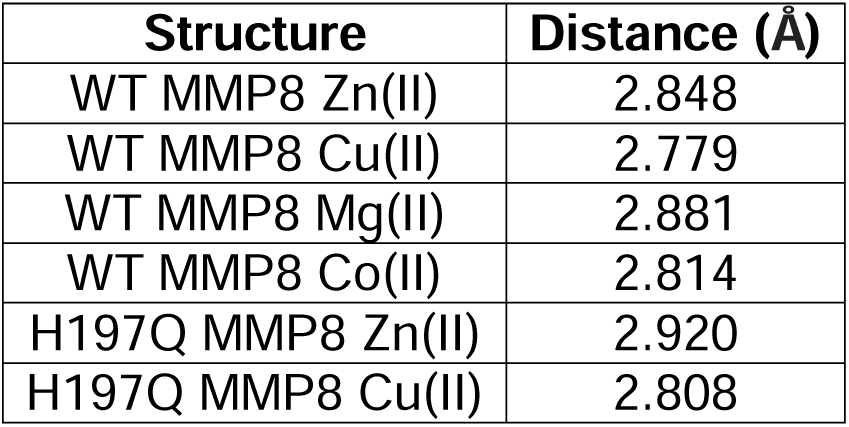
Relative distance between the water oxygen atom and the carbonyl carbon atom.

**Table S2.**
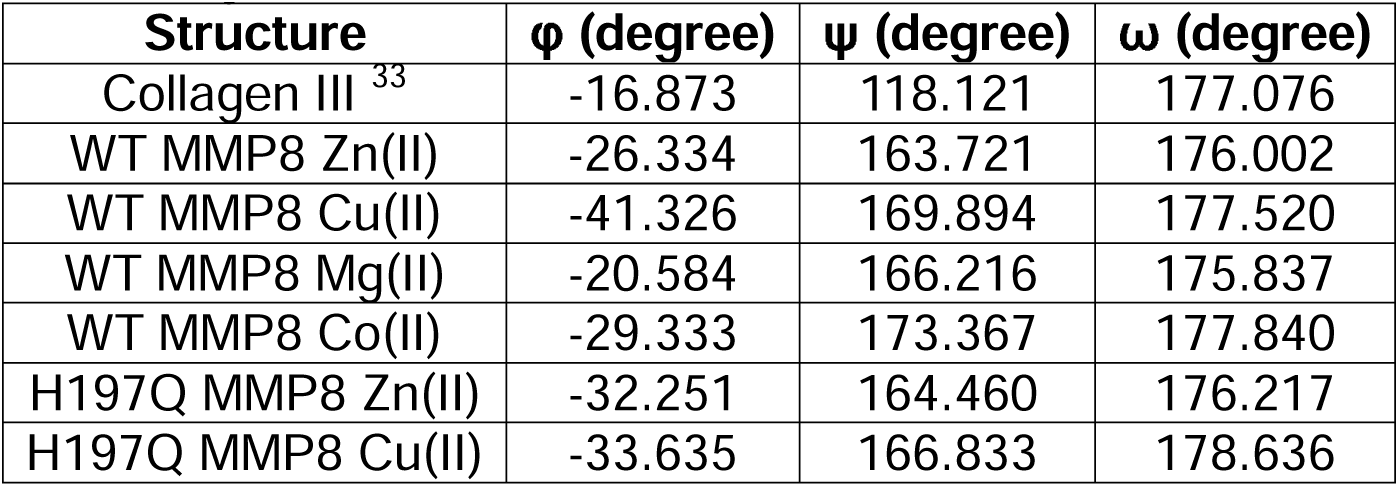
Torsion angles in the simulated ligand for the i position where peptide bond gets cleaved by MMP8.

## Conclusions

Computational analysis provides insights into the behaviors of MMP8 protein in the presence of different metal cofactors. In particular, the coordination geometries of the Zn(II) and Cu(II) occupied MMP8 are predicted and examined. It is found that Jhan-Teller effects destabilizes the Cu(II) occupied MMP8 by changing ligand binding from a bi-dentate to a mono-dentate fashion. In addition, simulations predict that H197Q is likely going to be able to utilize Cu(II) in catalytic reactions. In simulations, favorable ligand torsion angles at the enzyme cleavage position are predicted, and the optimized angles were found to differ from those measured in the crystal structure of collagen ligand, suggesting that a ligand conformational change is required before MMP8 cleavage can take place. The insights provided by these simulations provides directions for future protein engineering efforts based on the MMP8 protein.

## References

1. Nagase, H.; Woessner, J. F., Jr., Matrix metalloproteinases. J Biol Chem 1999, 274 (31), 21491–4.

2. Kleiner, D. E., Jr.; Stetler-Stevenson, W. G., Structural biochemistry and activation of matrix metalloproteases. Curr Opin Cell Biol 1993, 5 (5), 891–7.

3. Overall, C. M.; Lopez-Otin, C., Strategies for MMP inhibition in cancer: innovations for the post-trial era. Nat Rev Cancer 2002, 2 (9), 657–72.

4. Fields, G. B., The Rebirth of Matrix Metalloproteinase Inhibitors: Moving Beyond the Dogma. Cells 2019, 8 (9).

5. Macartney, H. W.; Tschesche, H., The metal ion requirement for activation of latent collagenase from human polymorphonuclear leucocytes. Hoppe Seylers Z Physiol Chem 1981, 362 (11), 1523–31.

6. Hahn, P. F.; Fairman, E., The copper content of some human and animal tissues. Journal of Biological Chemistry 1936, 113 (1), 161–165.

7. Margalioth, E. J.; Schenker, J. G.; Chevion, M., Copper and zinc levels in normal and malignant tissues. Cancer 1983, 52 (5), 868–72.

8. Rubino, J. T.; Franz, K. J., Coordination chemistry of copper proteins: How nature handles a toxic cargo for essential function. J Inorg Biochem 2012, 107 (1), 129–143.

9. Ye, S.; Wu, X.; Wei, L.; Tang, D. M.; Sun, P.; Bartlam, M.; Rao, Z. H., An insight into the mechanism of human cysteine dioxygenase - Key roles of the thioether-bonded tyrosine-cysteine cofactor. Journal of Biological Chemistry 2007, 282 (5), 3391–3402.

10. Lentsch, A. B.; Kato, A.; Saari, J. T.; Schuschke, D. A., Augmented metalloproteinase activity and acute lung injury in copper-deficient rats. Am J Physiol Lung Cell Mol Physiol 2001, 281 (2), L387–93.

11. Parr-Sturgess, C. A.; Tinker, C. L.; Hart, C. A.; Brown, M. D.; Clarke, N. W.; Parkin, E. T., Copper modulates zinc metalloproteinase-dependent ectodomain shedding of key signaling and adhesion proteins and promotes the invasion of prostate cancer epithelial cells. Mol Cancer Res 2012, 10 (10), 1282–93.

12. Hwang, J. J.; Park, M. H.; Koh, J. Y., Copper activates TrkB in cortical neurons in a metalloproteinase-dependent manner. J Neurosci Res 2007, 85 (10), 2160–6.

13. Heitzer, M.; Hallmann, A., An extracellular matrix-localized metalloproteinase with an exceptional QEXXH metal binding site prefers copper for catalytic activity. J Biol Chem 2002, 277 (31), 28280–28286.

14. McGuffin, L. J.; Adiyaman, R.; Maghrabi, A. H.; Shuid, A. N. A.; Brackenridge, D. A.; Nealon, J. O.; Philomina, L. S., IntFOLD: an integrated web resource for high performance protein structure and function prediction. Nucleic Acids Res 2019, 47 (W1), W408–W413.

15. Juurikka, K.; Butler, G. S.; Salo, T.; Nyberg, P.; Astrom, P., The Role of MMP8 in Cancer: A Systematic Review. Int J Mol Sci 2019, 20 (18).

16. Coates, R. J.; Weiss, N. S.; Daling, J. R.; Rettmer, R. L.; Warnick, G. R., Cancer risk in relation to serum copper levels. Cancer Res 1989, 49 (15), 4353–6.

17. Denoyer, D.; Masaldan, S.; La Fontaine, S.; Cater, M. A., Targeting copper in cancer therapy: ‘Copper That Cancer’. Metallomics 2015, 7 (11), 1459–76.

18. Bode, W.; Gomis-Ruth, F. X.; Stockler, W., Astacins, serralysins, snake venom and matrix metalloproteinases exhibit identical zinc-binding environments (HEXXHXXGXXH and Met-turn) and topologies and should be grouped into a common family, the ‘metzincins’. FEBS Lett 1993, 331 (1-2), 134–40.

19. Tauro, M.; Laghezza, A.; Loiodice, F.; Piemontese, L.; Caradonna, A.; Capelli, D.; Montanari, R.; Pochetti, G.; Di Pizio, A.; Agamennone, M.; Campestre, C.; Tortorella, P., Catechol-based matrix metalloproteinase inhibitors with additional antioxidative activity. J Enzyme Inhib Med Chem 2016, 31 (sup4), 25–37.

20. Bertini, I.; Calderone, V.; Fragai, M.; Luchinat, C.; Maletta, M.; Yeo, K. J., Snapshots of the reaction mechanism of matrix metalloproteinases. Angew Chem Int Ed Engl 2006, 45 (47), 7952–5.

21. Pelmenschikov, V.; Siegbahn, P. E., Catalytic mechanism of matrix metalloproteinases: two-layered ONIOM study. Inorg Chem 2002, 41 (22), 5659–66.

22. Lovejoy, B.; Hassell, A. M.; Luther, M. A.; Weigl, D.; Jordan, S. R., Crystal-Structures of Recombinant 19-Kda Human Fibroblast Collagenase Complexed to Itself. Biochemistry-Us 1994, 33 (27), 8207–8217.

23. Stocker, W.; Bode, W., Structural Features of a Superfamily of Zinc-Endopeptidases - the Metzincins. Curr Opin Struc Biol 1995, 5 (3), 383–390.

24. Manka, S. W.; Carafoli, F.; Visse, R.; Bihan, D.; Raynal, N.; Farndale, R. W.; Murphy, G.; Enghild, J. J.; Hohenester, E.; Nagase, H., Structural insights into triple-helical collagen cleavage by matrix metalloproteinase 1. Proc Natl Acad Sci U S A 2012, 109 (31), 12461–6.

25. Prior, S. H.; Byrne, T. S.; Tokmina-Roszyk, D.; Fields, G. B.; Van Doren, S. R., Path to Collagenolysis: COLLAGEN V TRIPLE-HELIX MODEL BOUND PRODUCTIVELY AND IN ENCOUNTERS BY MATRIX METALLOPROTEINASE-12. J Biol Chem 2016, 291 (15), 7888–901.

26. Koch, K. A.; Pena, M. M.; Thiele, D. J., Copper-binding motifs in catalysis, transport, detoxification and signaling. Chem Biol 1997, 4 (8), 549–60.

27. Warren, J. J.; Lancaster, K. M.; Richards, J. H.; Gray, H. B., Inner- and outer-sphere metal coordination in blue copper proteins. J Inorg Biochem 2012, 115, 119–26.

28. Nar, H.; Huber, R.; Messerschmidt, A.; Filippou, A. C.; Barth, M.; Jaquinod, M.; van de Kamp, M.; Canters, G. W., Characterization and crystal structure of zinc azurin, a by-product of heterologous expression in Escherichia coli of Pseudomonas aeruginosa copper azurin. Eur J Biochem 1992, 205 (3), 1123–9.

29. Stiebritz, M. T.; Hu, Y., Computational Methods for Modeling Metalloproteins. Methods Mol Biol 2019, 1876, 245–266.

30. Gogoi, P.; Chandravanshi, M.; Mandal, S. K.; Srivastava, A.; Kanaujia, S. P., Heterogeneous behavior of metalloproteins toward metal ion binding and selectivity: insights from molecular dynamics studies. J Biomol Struct Dyn 2016, 34 (7), 1470–85.

31. Siegbahn, P. E.; Himo, F., Recent developments of the quantum chemical cluster approach for modeling enzyme reactions. J Biol Inorg Chem 2009, 14 (5), 643–51.

32. Borowski, T.; Quesne, M.; Szaleniec, M., QM and QM/MM Methods Compared: Case Studies on Reaction Mechanisms of Metalloenzymes. Adv Protein Chem Struct Biol 2015, 100, 187–224.

33. Boudko, S. P.; Engel, J.; Okuyama, K.; Mizuno, K.; Bachinger, H. P.; Schumacher, M. A., Crystal structure of human type III collagen Gly991-Gly1032 cystine knot-containing peptide shows both 7/2 and 10/3 triple helical symmetries. J Biol Chem 2008, 283 (47), 32580–9.

34. Tochowicz, A.; Maskos, K.; Huber, R.; Oltenfreiter, R.; Dive, V.; Yiotakis, A.; Zanda, M.; Pourmotabbed, T.; Bode, W.; Goettig, P., Crystal structures of MMP-9 complexes with five inhibitors: contribution of the flexible Arg424 side-chain to selectivity. J Mol Biol 2007, 371 (4), 989–1006.

35. Sakharov, D. V.; Lim, C., Zn protein simulations including charge transfer and local polarization effects. J Am Chem Soc 2005, 127 (13), 4921–9.

