## Supplemental Figure and Tables for "Computational Analysis of the Metal Selectivity of Matrix Metalloproteinase 8"

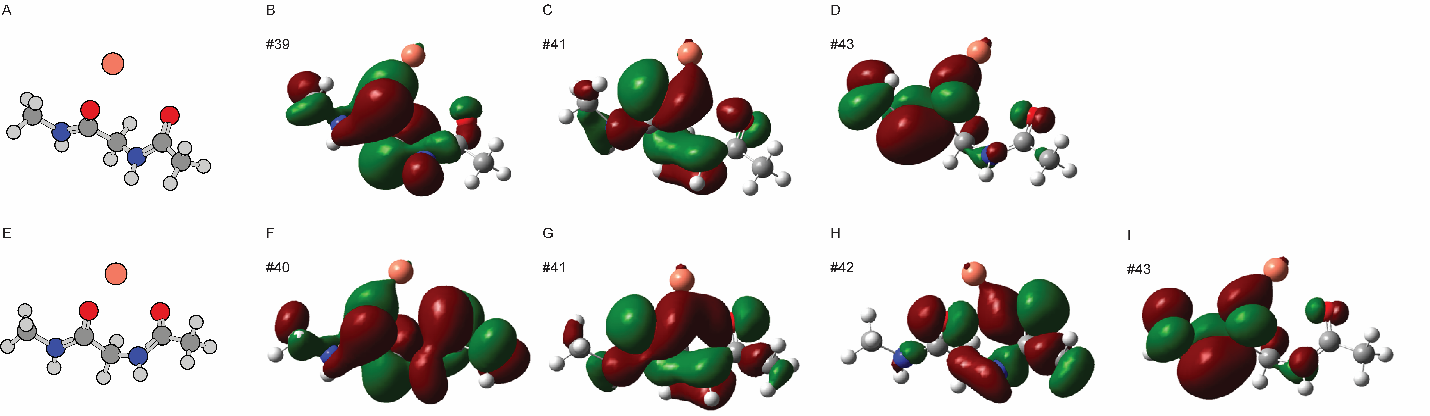

**Figure S1.** Cu(II) –ligand orbital mixing the bonding orbitals formed between the ligand and Cu(II) ion are shown. (A) is a molecular representation of the Cu(II)-ligand complex in wild-type MMP8 catalytic site. (B)-(D) are three bonding molecular orbitals between Cu(II) and ligand wild-type MMP8 catalytic site. (E) is a molecular representation of the Cu(II)-ligand complex in H197Q mutant catalytic site. (B)-(D) are four bonding molecular orbitals between Cu(II) and ligand in H197Q mutant catalytic site.

**Table S1.** Relative distance between the water oxygen atom and the carbonyl carbon atom.

| **Structure** | **Distance (Å)** |
| --- | --- |
| WT MMP8 Zn(II) | 2.848 |
| WT MMP8 Cu(II) | 2.779 |
| WT MMP8 Mg(II) | 2.881 |
| WT MMP8 Co(II) | 2.814 |
| H197Q MMP8 Zn(II) | 2.920 |
| H197Q MMP8 Cu(II) | 2.808 |

**Table S2.** Torsion angles in the simulated ligand for the i position where peptide bond gets cleaved by MMP8.

| **Structure** | **φ (degree)** | **ψ (degree)** | **ω (degree)** |
| --- | --- | --- | --- |
| Collagen III ^33^ | -16.873 | 118.121 | 177.076 |
| WT MMP8 Zn(II) | -26.334 | 163.721 | 176.002 |
| WT MMP8 Cu(II) | -41.326 | 169.894 | 177.520 |
| WT MMP8 Mg(II) | -20.584 | 166.216 | 175.837 |
| WT MMP8 Co(II) | -29.333 | 173.367 | 177.840 |
| H197Q MMP8 Zn(II) | -32.251 | 164.460 | 176.217 |
| H197Q MMP8 Cu(II) | -33.635 | 166.833 | 178.636 |

**References**

33.	Boudko, S. P.; Engel, J.; Okuyama, K.; Mizuno, K.; Bachinger, H. P.; Schumacher, M. A., Crystal structure of human type III collagen Gly991-Gly1032 cystine knot-containing peptide shows both 7/2 and 10/3 triple helical symmetries. *J Biol Chem* **2008,** *283* (47), 32580-9.
